# Robust phylogenetic regression

**DOI:** 10.1101/2022.08.26.505424

**Authors:** Richard Adams, Zoe Cain, Raquel Assis, Michael DeGiorgio

## Abstract

Modern comparative biology owes much to phylogenetic regression. At its conception, this technique sparked a revolution that armed biologists with phylogenetic comparative methods (PCMs) for combatting evolutionary pseudoreplication, which arises inherently from trait data sampled across related species. Over the past few decades, the phylogenetic regression framework has become a paradigm of modern comparative biology that has been widely embraced as a remedy for evolutionary pseudoreplication. However, recent evidence has sown doubt over the efficacy of phylogenetic regression, and PCMs more generally, with the suggestion that many of these methods fail to provide an adequate defense against unreplicated evolution—the primary justification for using them in the first place. Importantly, some of the most compelling examples of biological innovation in nature result from abrupt, lineage-specific evolutionary shifts, which current regression models are largely ill-equipped to deal with. Here we explore a solution to this problem by applying robust linear regression to comparative trait data. We formally introduce robust phylogenetic regression to the PCM toolkit with linear estimators that are less sensitive to model violations while still retaining high power to detect true trait associations. Our analyses also highlight an ingenuity of the original algorithm for phylogenetic regression based on independent contrasts, whereby robust estimators are particularly effective. Collectively, we find that robust estimators hold promise for improving tests of trait associations and offer a path forward in scenarios where classical approaches may fail. Our study joins recent arguments for increased vigilance against pseudoreplication and a better understanding of evolutionary model performance in challenging–yet biologically important–settings.

## Introduction

Since Darwin’s time, biologists have struggled to understand the evolutionary dynamics among organisms and their traits that have collectively shaped present-day biodiversity. One of the most notorious sources of headaches for modern comparative biologists is evolutionary pseudoreplication (Fig. 1), a widespread phenomenon in which measured traits tend to covary according to the hierarchical structure of the phylogenetic tree relating the species in which they were sampled (Felsenstein 1985; Grafen 1989; Martins and Hansen 1997; Pagel 1997, 1999; Rohlf 2001). Consequently, related species and their traits represent pseudoreplicates rather than independent observations, violating a common assumption of statistical tests. Failure to account for evolutionary pseudoreplication can therefore bias inferences toward incorrect conclusions with high confidence (Felsenstein 1985; Maddison and FitzJohn 2015; Uyeda et al. 2018), such that it is now broadly acknowledged that one must consider the phylogenetic background upon which organisms and their traits have evolved.

**FIGURE 1.**
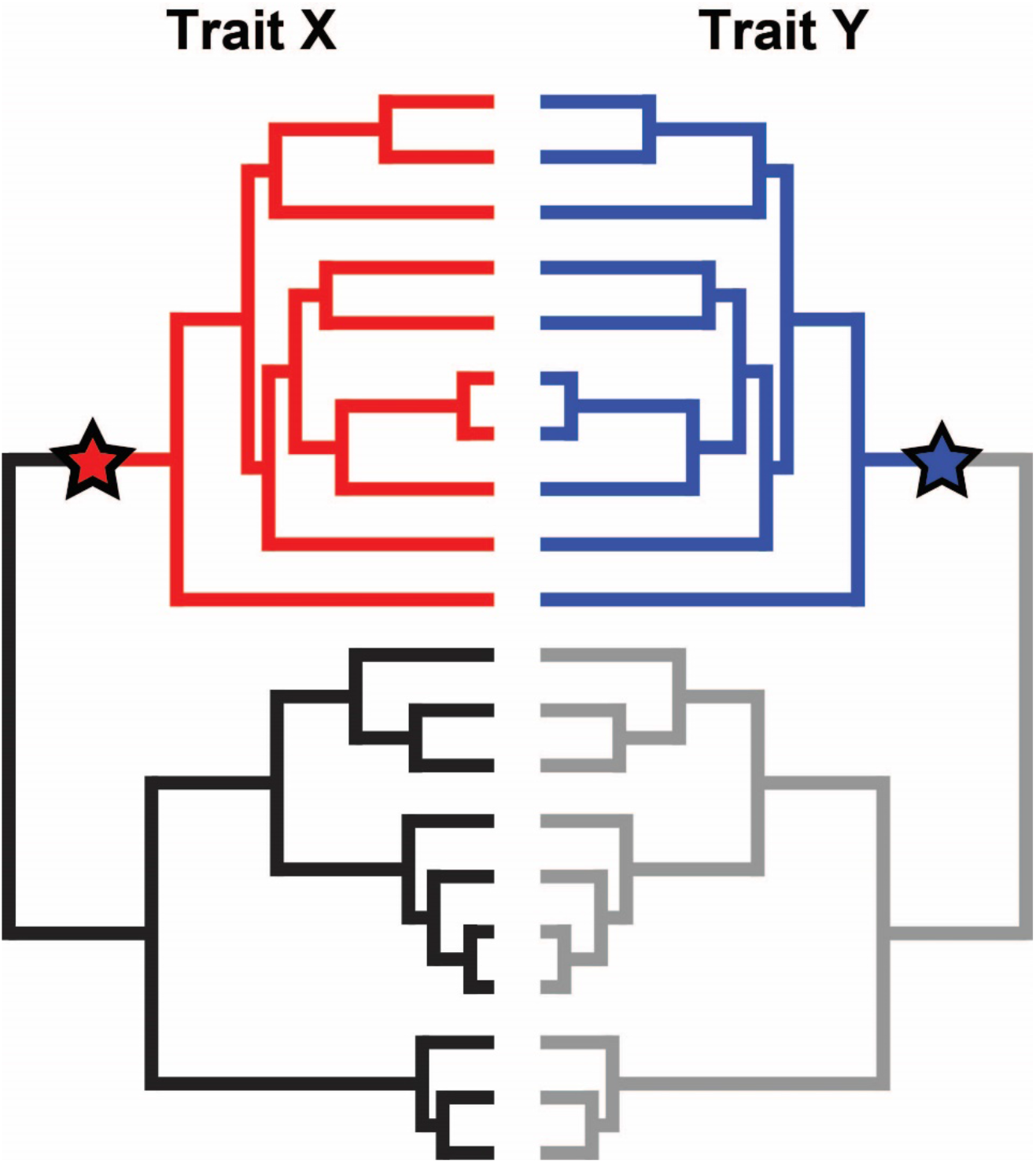
Evolutionary pseudoreplication coupled with independent trait shifts can lead to false associations between traits. In this example, observed values for traits X and Y are not independent across species (i.e., the species are pseudoreplicates) due to the hierarchical structure of the sampled species. A single shift in the trait distribution (marked by stars) affects a large number of species due to their shared ancestry, leading to a false signal of correlated evolution if not properly accounted for. Figure inspired by Figure 2 of Uyeda et al. (2018).

Nearly 40 years ago, Felsenstein (1985) proposed a simple yet elegant solution to this problem: phylogenetic regression, in which the statistical model is informed by tree structure. In doing so, he established phylogenetic comparative methods (PCMs) as the *de facto* standard in the field, ushering in a new era of tree thinking in comparative biology (Carvalho et al. 2005; Huey et al. 2019). The principles of phylogenetic regression were further clarified and expanded upon with the application of generalized least squares (Grafen 1989; Martins and Hansen 1997; Pagel 1997, 1999; Rohlf 2001), which accounts for statistical dependence by explicitly modeling the correlated error structure that results from shared inheritance. Over the past few decades, phylogenetic regression has become a paradigm of modern comparative biology, inspiring a wealth and diversity of offspring approaches for studying a myriad of biological hypotheses and questions under the PCM umbrella (e.g., Harvey and Pagel 1991; Blomberg et al. 2003; Felsenstein 2004; O’Meara et al. 2006; Revell et al. 2008; Beaulieu et al. 2012; Pennell and Harmon 2013). Importantly, a major goal of phylogenetic regression, and PCMs more generally, is to disentangle evidence of adaptation from simple inheritance, and thus, these methods have become widely embraced as a remedy for evolutionary pseudoreplication.

However, recent studies have sown doubt over the cannon of phylogenetic regression, with evidence suggesting that PCMs may not be the panacea as long hoped (Maddison and FitzJohn 2015; Uyeda et al. 2018). Ironically, many of these methods seemingly fail to protect against evolutionary pseudoreplication—the reason they were developed in the first place. Deeper introspection into the justification and philosophical underpinnings of PCMs led to a recent call for a complete “rethinking” of the current paradigm (Uyeda et al. 2018), suggesting that biologists may be seriously overestimating evidence of trait associations (Maddison and FitzJohn 2015). At the heart of these concerns is the realization that widely held assumptions about trait evolution may not always reflect reality. In particular, a fundamental assumption of classical phylogenetic regression is that evolution proceeds more or less as a continuous process that can be approximated using Brownian motion (BM; Cavalli-Sforza and Edwards 1967; Felsenstein 1973) or related models that extend BM principles (Lande 1979; Hansen 1997; Pagel 1999; Blomberg et al. 2003; Harmon et al. 2010). Yet a breadth of macroevolutionary data suggests that biodiversity has been profoundly shaped by abrupt evolutionary shifts and discontinuities that often act in a lineage-specific manner, violating core PCM assumptions (Fig. 1; Schluter 2000; Uyeda et al. 2011, 2017; Slater and Pennell 2014; Landis and Schraiber 2017).

Scenarios of “evolution by jumps” were first hypothesized by Simpson (1944) to describe rapid and dramatic phenotypic shifts in response to changes in the adaptive landscape (e.g., new environments, empty niches, and key innovations), which often manifest as lineage-specific novelties. Whereas more realistic models of trait shifts have been proposed in several contexts (Eastman et al. 2011; Bartoszek et al. 2012; Uyeda and Harmon 2014; Clavel et al. 2015; Bastide et al. 2018), phylogenetic regression is typically implemented under the assumption of continuous trait change in the absence of such phenomena. Current techniques are thus ill-equipped to deal with the dynamics observed in nature (O’Meara 2012; Pennell and Harmon 2013; Garamszegi 2014; Mazel et al. 2016), yielding systematic error that results from the inability of models to distinguish true statistical associations from instantaneous evolutionary shifts (Maddison and FitzJohn 2015; Uyeda et al. 2018).

In this study, we explore a new solution to this problem: the application of robust regression (Huber 2004; Yu and Yao 2017) for testing statistical trait associations. Mirroring the history of PCMs, “robust statistics” emerged during the 1980s as a new branch of statistics born out of concern for the rigor of classic techniques in the presence of model violations and outliers (Huber 2004; Yu and Yao 2017). Yet, save for a handful of examples (Slater and Pennell 2014; Arbour and López-Fernández 2016; Puttick 2018), robust methods have been largely overlooked in comparative trait studies, such that there has been an almost singular focus on using classical regression models to infer trait associations. Here we introduce robust phylogenetic regression to the PCM toolkit with four linear estimators, and we evaluate their performances across an array of statistically challenging and yet biologically important scenarios of rapid, lineage-specific evolutionary shifts known to mislead traditional phylogenetic regression. Following the protocol of a previous study that revealed biases in classical PCMs under such conditions (Uyeda et al. 2018), we examine the application of robust phylogenetic regression to traits that have experienced episodes of instantaneous jumps in trait space. To investigate the properties of these estimators for phylogenetic regression, we probe their behavioral characteristics on an array of simulated and empirical examples.

## Methods

### The Linear Model, Comparative Trait Data, and Phylogenetic Regression

Linear regression is arguably the most commonly applied statistical technique in biology (Sokal and Rohlf 1981; Ford 2000; Queen et al. 2002), as well as in the sciences more generally (Montgomery et al. 2012; Seber and Lee 2012), for studying relationships between variables. The familiar linear regression equation can be written as

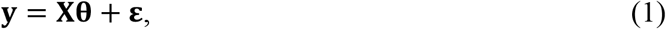

where 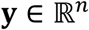 is an *n*-dimensional vector of the response variable, 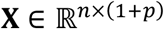 is an *n* × (1 +*p*) design matrix (first column is a vector 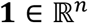 with the value of one for each element) containing *n* measurements for each of *p* predictor variables, 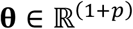 denotes the vector of unknown model parameters, and 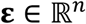 represents the residuals, or errors in predicting the response variable **y**. Given a dataset 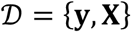, linear regression seeks to approximate **θ** by finding the optimal estimates 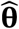 that minimize a cost function summarizing the overall magnitude of the residuals **ε** = **y** – **ŷ**, which are computed as the difference between the observed response values **y** and the model predictions 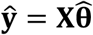. Importantly, accuracy of the estimates 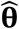 and associated tests of significance (Rencher and Schaalje 2008) depend on fundamental assumptions of the model (Poole and O’Farrell 1971; Montgomery et al. 2012; Seber and Lee 2012; Mundry 2014). One such assumption of ordinary least squares (OLS) regression is that the errors **ε** are independently and identically distributed (i.i.d) as normal, such that 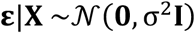 follows a multivariate normal distribution, where 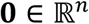 is an *n*-dimensional column vector representing the expected value of zero for each element in **ε**, 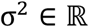 specifies a constant and unknown variance, and 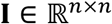 is the *n* × *n* identity matrix.

Phylogenetic regression strives to fit Equation 1 to test for a statistical relationship between a response trait **y** and one or more predictor traits **X** measured in 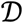 while explicitly considering the phylogenetic background upon which they evolved. Accounting for phylogeny requires correcting for the assumption of OLS that measurements of the error terms **ε** are uncorrelated. Researchers face two options for this task: phylogenetic independent contrasts (PIC; Felsenstein 1985) and phylogenetic generalized least squares (PGLS; Grafen 1989; Martins and Hansen 1997; Pagel 1997, 1999; Rohlf 2001). Both PIC and PGLS recognize that traits measured across related species are not statistically independent and use different strategies to address this issue. PIC computes a series of statistically independent contrasts according to the algorithm of Felsenstein (1985), yielding a newly transformed dataset 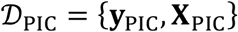, where 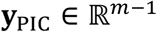 and 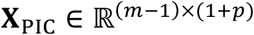 compose the collection of *m* – 1 contrasts in *m* species for the response trait **y** and predictor traits **X**, respectively.

PGLS expands upon PIC by explicitly modeling a covariance structure **C** into the residual error, such that 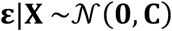, where 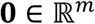 and 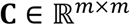 is the *m* × *m* phylogenetic covariance matrix (Felsenstein 1973; O’Meara et al. 2006) specified according to a particular tree and assumed evolutionary model (Grafen 1989; Martins and Hansen 1997; Pagel 1997, 1999; Rohlf 2001). Following the established OLS transformation of GLS (Judge and Griffiths 1985; Kariya and Kurata 2004; Rencher and Schaalje 2008), PGLS can also be implemented as a projection of traits through a phylogenetic transformation matrix 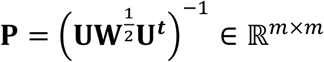, where the eigenvectors 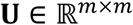 and *m* associated eigenvalues along diagonal of diagonal matrix 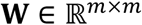 are obtained from eigendecomposition of **C** = **UWU**^−1^(Garland Theodore and Ives 2000; Adams 2014; Adams and Collyer 2018). The original trait values can then be transformed by **P** to yield 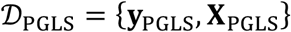, where **y**_PGLS_ = **Py** and **X**_PGLS_ = **PX**. By rescaling branch lengths based on trait evolution, modeling the residual error as a function of **C** allows flexibility to the extent of “phylogenetic signal” present in the data (Grafen 1989; Mundry 2014).

PGLS estimates are identical to those of classical OLS in the absence of signal, while phylogenetic covariance is corrected to an appropriate degree with intermediate signal (Grafen 1989; Mundry 2014). PIC can also adapt these considerations (Felsenstein 1985; Symonds and Blomberg 2014), though this strategy is less common in practice. Importantly, PIC and PGLS regression estimates, or coefficients, are equivalent under BM evolution (Grafen 1989; Garland Theodore and Ives 2000; Rohlf 2001; Blomberg et al. 2012; Symonds and Blomberg 2014). PGLS uses the phylogenetic covariance matrix to project the trait data, such that its coefficients can be visualized alongside the original trait measurements (Blomberg et al. 2012; Symonds and Blomberg 2014).

### The Ubiquitous Least Squares Estimator

Almost without exception, PIC (i.e., using 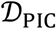) and PGLS (i.e., using 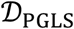) regression are conducted using the classical least squares (L2) estimator. The L2 estimator can be defined as

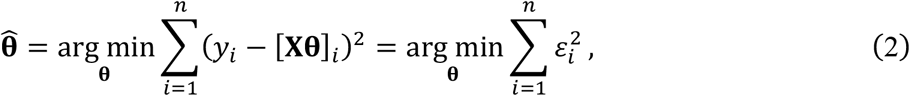

where *y_i_* denotes the known response and [**Xθ**]_*i*_ the predicted response given parameter vector **θ** of the *i*th contrast of *n* = *m* – 1 total contrasts for PIC regression, or of the *i*th species of *n* = *m* total species for PGLS regression, with *ε_i_* = *y_i_* – [**Xθ**]_*i*_ therefore measuring the *i*th residual for both PIC and PGLS regression. Equation 2 also leads to the closed-form solution for the parameter estimates 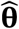 according to the normal equations 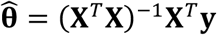, where **X**^*T*^ is the transpose of **X**, which can represent either **X**_PIC_ or **X**_PGLS_, and the −1 superscript indicates the matrix inverse. The L2 estimator makes no assumptions about the validity of the linear model (i.e., Eq. 1); it simply finds the regression coefficients that provide the best linear fit to the data by minimization with Equation 2. Another way of viewing the L2 estimator is that it minimizes the squared Euclidian distance, or *ℓ*_2_-norm, between the known values of **y** and the model predictions **ŷ**. The *ℓ_k_*-norm of a vector **v** is defined as 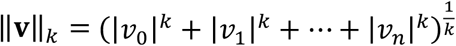 for finite integer *k* > 0. In general, higher indices *k* place more emphasis on larger values in **v**.

### Robust Phylogenetic Regression

Here we introduce “robust phylogenetic regression” to the PCM toolkit with the linear M, L1 (least absolute deviation), S, and MM estimators. It is well-established that the standard L2 estimator is highly sensitive to outliers, and these estimators replace least squares with robust criteria (Yu and Yao 2017). Each robust estimator therefore has favorable properties related to its breakdown point, or proportion of outliers that can be tolerated, and efficiency relative to the L2 estimator in the absence of outliers (Donoho and Huber 1983; Yu and Yao 2017). The L2 estimator has a breakdown point of 1/*n*, which tends to zero as the sample size *n* increases, meaning that even a single unusual observation can exert strong influence (Rousseeuw and Yohai 1984). Thus, robust estimators strive to achieve high breakdown point, efficiency, or both.

M estimators are a class of estimators representing solutions to the normal equations that incorporate appropriate weighting functions (Huber 1973, 1992), which have been used to detect early bursts of trait evolution in the presence of outlier taxa with posterior predictive model checks (Slater and Pennell 2014; Arbour and López-Fernández 2016). M estimators can be viewed as a form of weighted regression that seeks to downweigh the influence of large residuals that may bias inferences (Yu and Yao 2017). An iteratively reweighted least squares procedure is used to estimate the weights as (Holland and Welsch 1977)

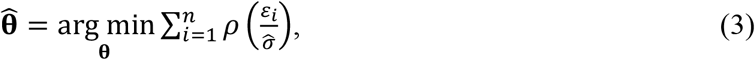

where *ρ*(*t*) is a robust loss function giving the weights of each residual *ε_i_*, and 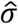 is an error scale estimate. The influence function *ρ*′(*t*) is the derivative of *ρ*(*t*), which is most commonly specified as Huber’s weighting scheme (Huber 1973, 2004), where *ρ′*(*t*) = max{−*c*, min(*c*, *t*)}, with *c* = 1.345 recommended in practice. Observations that do not deviate substantially from model predictions are effectively applied weights of one, and conversely, larger residuals are downweighed in the process. In particular, the solution to the L2 estimator is found when *ρ*(*t*) = *t*^2^.

We can also obtain the L1 estimator by setting *ρ*(*t*) = |*t*|, which minimizes the sum of the absolute values of the residuals as

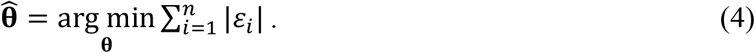

The L1 estimator minimizes the *ℓ*_1_-norm between the observed and predicted responses, which is designed to be more robust than the L2 estimator because Equation 4 minimizes the sum of absolute residual values |*ε_i_*|, rather than the sum of squared residuals 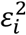, as in Equation 2. Hence, the L1 estimator de-emphasizes outlier residuals that may lead to large squared distances with high leverage, or unusual values of the predictor traits **X** (Rousseeuw and Yohai 1984), which have also been shown to bias classical L2-based phylogenetic regression (Uyeda et al. 2018). However, like the L2 estimator, the L1 estimator has a breakdown point of 1/*n*, which tends toward zero as *n* increases, and thus can be sensitive to high-leverage outliers (Maronna et al. 2019). In general, M estimators with monotone influence functions have a breakdown point of 1/*n* → 0 as *n* → ∞, which results in a lack of immunity to large outliers (Maronna et al. 2019).

S estimators (Rousseeuw and Yohai 1984) seek to minimize the dispersion of the residuals defined by

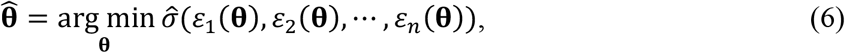

where *ε_i_*(**θ**) = *ε_i_* and 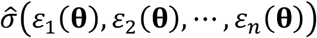 is the scaled M estimator defined as the solution to

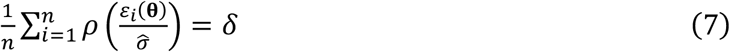

for any given values of parameters **θ**, where *δ* is taken to be *E*[*ρ*(***ε***)]. S estimators can achieve a high breakdown point of 0.5 with an asymptotic efficiency of 0.29 (Maronna et al. 2019).

MM estimators were first proposed by Yohai (1987), and now represent one of the most popular robust regression techniques (Yu and Yao 2017). MM estimation involves a three-step procedure: (1) an initial estimate 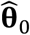 is obtained using an estimator with high breakpoint but potentially low efficiency, (2) a robust M estimate of the scale 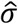 of the residuals is obtained from 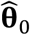, and (3) a final M estimate 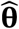 is found using 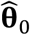 as a starting point (Yu and Yao 2017). In practice, the initial estimate can be obtained via an S estimator, followed by two successive rounds of M estimation to estimate 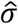 and the final 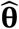. Typically, an MM estimate with a bisquare function (initial S estimate also starting from a bisquare function; Gross 1977) and efficiency 0.85 is recommended in practice (Yu and Yao 2017), which can yield an asymptotic breakdown point of 0.5.

For our analyses, we leveraged a rich library of robust regression software packages within the R statistical environment (R Core Team 2013). Specifically, we used the *rlm* function with *method*=“*M*” in the MASS package (Ripley 2015) for M estimation with iterated re-weighted least squares and Huber’s weighting scheme, the Zαd function in the L1PACK package (Osorio et al. 2017) for L1 estimation, the *lmRob* function with option *estim*=“*Initial*” in the ROBUST package (Maechler 2014) for S estimation, and the *lmRob* function in the ROBUST package for MM estimation. Classical L2 estimation was conducted using the *lm* function in base R. These functions have been wrapped into a new open-source R package, ROBRT (ROBust Regression on Trees), for conducting robust regression within a coordinated framework for studies of comparative trait evolution.

### Simulations With Evolutionary Shifts

We sought to understand the performance of robust regression for challenging–yet biological important–scenarios of rapid and unreplicated evolutionary change known to bias inferences (Uyeda et al. 2018). To do so, we first compared performances of the four robust estimators and the classical L2 estimator on datasets simulated across a wide range of increasing model misspecification severity. In these scenarios, we set the true regression coefficients in **θ** to zero to represent uncorrelated (statistically independent) trait evolution, allowing us to evaluate the false positive rate for detecting correlated trait evolution of each estimator.

Specifically, we generated bivariate datasets 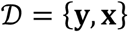 containing measurements of two statistically independent traits 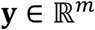 and 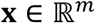 simulated under a model of evolutionary shifts (termed “shift” model) for a phylogeny of *m* species (Figs. 2a and b). In our shift model, we assumed that **y** and **x** shifted simultaneously, but with differing magnitudes, at the same time and location (node in the tree), while all other parameters remained stationary throughout the tree (Eastman et al. 2013). We modified the classical BM process (Felsenstein 1973) to model the evolution of two independent traits by incorporating a pair of shift magnitudes 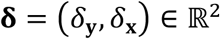 with standard evolutionary variances corresponding to BM rates 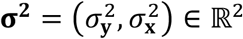 (O’Meara et al. 2006; Revell 2008) for traits **y** and **x**. The shift magnitudes **δ** were applied at the beginning of a single branch directly subtending the root that splits the tree into two clades of equal size (vertical dashes in Figs. 2a and b), which is predicted to yield high-leverage outliers that bias phylogenetic regression (Uyeda et al. 2018). To model variability in the shift magnitudes, we randomly sampled values of **δ** across simulations from a bivariate normal distribution with zero mean, zero covariance (independent *δ*_y_ and *δ*_x_), and variances that are scalar multiples of the BM rates **σ^2^**. That is, the simulated shift magnitudes are distributed as 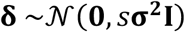, where **0** = (0,0), and *s***σ^2^I** represents the product of a scalar variance *s* > 0, the BM rates **σ^2^**, and the 2 × 2 identity matrix **I**. Throughout our simulations, we used a standard BM rate of one for each trait by setting **σ^2^** = (1,1) and the function *sim.char* from the GEIGER (Pennell et al. 2014) R package to simulate the ancestral (pre-shift) trait values with a mean state of zero for both **y** and **x**. Importantly, because traits **y** and **x** were independent under this shift model, the true slope parameter relating them was always zero.

**FIGURE 2.**
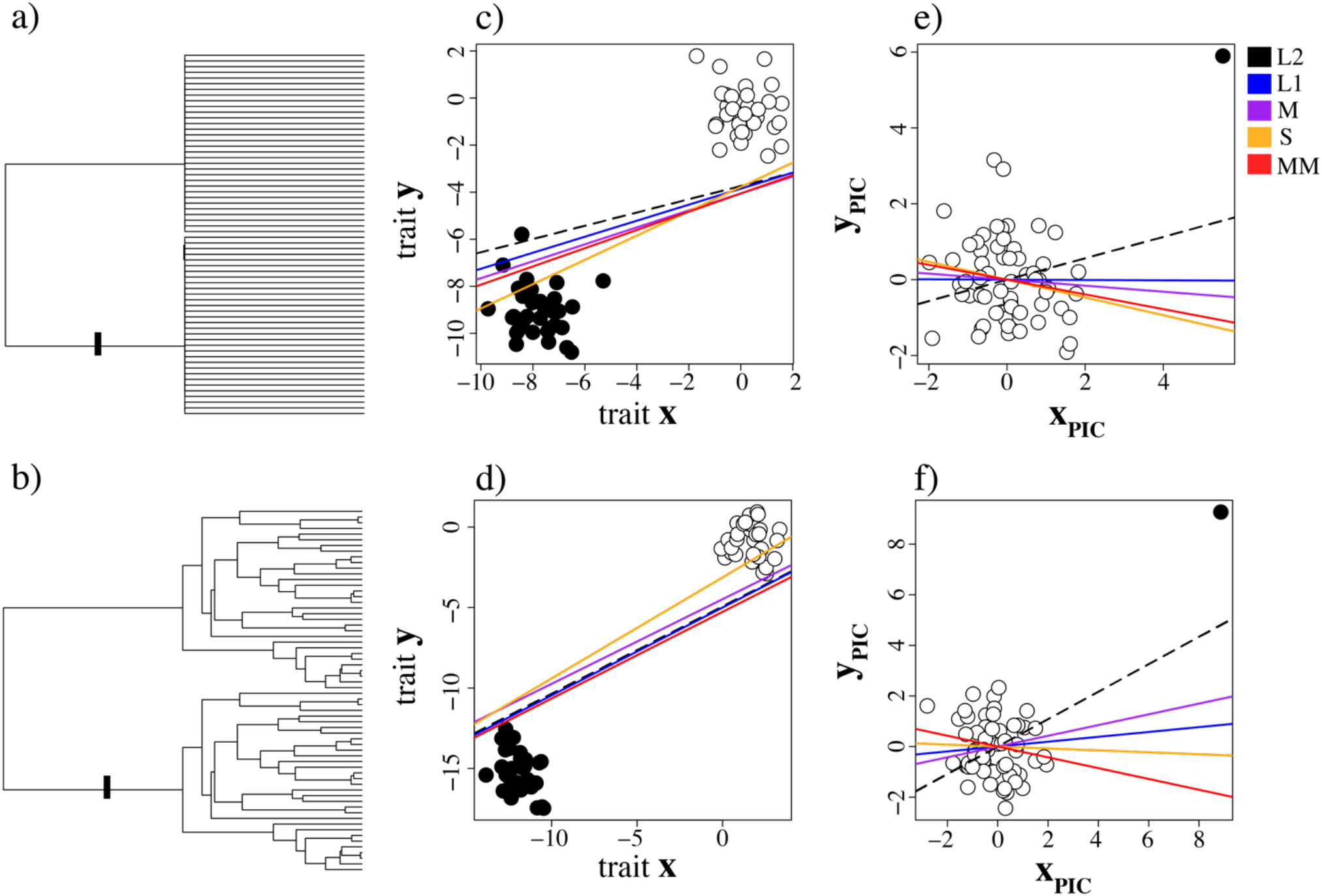
Characterizing differences among five linear estimates (L2, M, L1, S, and MM) for phylogenetic regression applied to statistically independent traits **y** and **x**. Data were simulated under the shift model with variance *s* = 10^2^ according to Felsenstein’s worst-case scenario (a) and a simulated balanced birth tree (b), with shift locations marked by dashes in corresponding trees. Covariation between traits **y** and **x** was assayed with PGLS (c and d) regression, and between transformed traits **y**_PIC_ and **x**_PIC_ with PIC regression (e and f). For graphical purposes, results shown in panels (c) and (d) are visualized in the original trait space–an advantage of PGLS.

Our simulation protocol generally followed the approach of Uyeda et al. (2018). First, we selected a shift variance *s* ∈ {10^−2^, 10^−1^,…, 10^5^}. Second, we used *s* to generate two independent values of the shift magnitudes 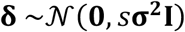. Third, we simulated a trait dataset 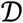 under a shift model with **δ**. Fourth, we transformed 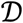 into 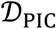 for PIC regression and 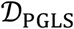 for PGLS regression. Last, we employed each of the five estimators (L2, M, L1, S, and MM) to perform PIC and PGLS regression on their transformed trait datasets. Given its popularity in comparative trait studies (Mazel et al. 2016), we used the default options for the classical *t* test with significance level *α* = 0.05 and *n* = *m* – 1 and *n* = *m* – 2 degrees of freedom for PIC and PGLS regression, respectively. We explored the performance of each estimator across diverse phylogenetic backgrounds using randomly generated trees of varying sizes corresponding to the number of species *m*. For each tree size *m* ∈ {64,128,256,512,1024}, we applied the scripts of Uyeda et al. (2018) to generate 100 balanced bifurcating trees by first simulating a pair of equally-sized subtrees with *m*/2 species under a pure birth model, and subsequently joining them together at the root (Figs. 2a and b). For each of the 100 trees, we simulated 100 replicate datasets according to the protocol described above for each value of *s* ∈ {10-^2^, 10^−1^,…, 10^5^}.

### Simulations With True Coevolutionary relationships

In addition to assaying false positive rates with simulations based on our shift model (Figs. 2a and b), we conducted a series of analyses to evaluate the statistical power (i.e., true positive rate) of each estimator for recovering a true coevolutionary relationship between two traits **y** and **x**. For these studies, we simulated trait data using a coevolutionary model (termed “coevo” model) of bivariate BM evolution (Revell and Harmon 2008) that varied the strength of the statistical association between the traits *ρ* ∈ {0.1,0.2,…, 1.0}. Specifically, we modeled the evolution of the traits **y** and **x** with among-trait covariance matrix

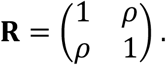

Each element of **R** corresponds to a particular evolutionary covariance term, with the first row and column corresponding to the response trait **y**, and the second row and column corresponding to the predictor trait **x**. The antidiagonal values *ρ* denote the degree of covariance between **y** and **x**, and the diagonal elements indicate BM rates of 1 for both **y** and **x** (i.e., 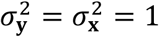).

We first selected the true among-trait covariance from *ρ* ∈ {0.1, 0.2,…, 1.0, 2.0,…, 5.0}. Next, we simulated traits **y** and **x** according to a bivariate BM model with among-trait covariance matrix **R** containing *ρ* and an ancestral state of zero. Last, we performed PIC and PGLS regression with each of the five estimators (L2, M, L1, S, and MM) to assess parameter estimates and significance of coevolution between traits **y** and **x**. Simulations under this coevo model did not include evolutionary shifts in the trait space, i.e., **δ** = (0,0). Mirroring the shift model simulations, we generated 100 balanced Yule trees using scripts provided by Uyeda et al. (2018), as well as 100 replicate datasets for each tree while varying the number of species *m* ∈ {64,128, 256, 512,1024}.

### Empirical Analyses

We applied robust phylogenetic regression to an empirical gene expression dataset from 11 female and male tissues in eight mammals and chicken (Brawand et al. 2011). Specifically, we obtained normalized gene expression abundance measurements computed in reads per kilobase of exon model per million mapped reads (RPKM; Mortazavi et al. 2008) from female and male brain (whole brain without cerebellum), female and male cerebellum, female and male heart, female and male kidney, female and male liver, and testis in human (*Homo sapiens*), chimpanzee (*Pan trogodytes*), gorilla (*Gorilla gorilla*), orangutan (*Pongo pygmaeus abelii*), macaque (*Macaca mulatta*), mouse (*Mus musculus*), opossum (*Monodelphis domestica*), platypus (*Ornithorhynchus anatinus*), and chicken (*Gallus gallus*; Brawand et al. 2011). We restricted our analysis to the most conservative 5,321 1:1 orthologs, or those with constitutive exons that aligned across all species in the dataset (Brawand et al. 2011), and computed the median expression level for tissues containing multiple replicates. Along with this expression dataset, we downloaded the gene expression phylogenetic tree constructed by Brawand et al. (2011), and scaled the tree depth to unit height. We investigated the statistical performance of the five estimators (L2, M, L1, S, and MM) in three experimental settings: expression in female brain ~ male brain, female heart ~ male heart, and female kidney ~ male kidney. For each experiment, we conducted PIC regression based on log-transformed RPKM values across the nine species, and explored relationships between conditions via evaluations of statistical significance (*P*-values) and estimated slope coefficients 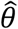.

## Results

### Demonstrating robust phylogenetic regression

Our simulations under the shift model (Fig. 2) represented opportunities to assess the sensitivity of each regression estimator (L2, M, S, and MM) to model violations because the two traits **y** and **x** were always statistically independent of one another. That is, the true slope coefficient of the linear model (Eq. 1) was *θ* = 0 in these scenarios, and our hope was that robust estimators would therefore fail to reject the null hypothesis of uncorrelated evolution by returning a non-significant *P*-value. We illustrated the application of robust phylogenetic regression in two familiar examples: Felsenstein’s worst case scenario (Fig. 2a) and a simple bifurcating tree (Fig. 2b). In both cases, all estimators applied to PGLS-transformed values displayed false positive trait associations (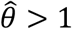; Figs. 2c and d), because species experiencing the ancestral trait shift (black circles) reside in a different space than those without the shift (white circles). In contrast, all robust estimators (M, L1, S, and MM) showed decreased sensitivity with PIC regression to the single high-leverage outlier (black circle; Figs. 2e and f), which represents the contrast generated by the branch with the ancestral trait shift, whereas the remaining contrasts cluster together (white circles). However, the L2 estimator is misled by this outlier contrast, exhibiting false positive trait associations (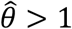; Figs. 2e and f). These examples hint that the application of robust regression estimators to PIC-transformed values may provide a solution to erroneous trait associations in the presence of such model violations.

### Robust Regression is Less Sensitive to Evolutionary Model Violations

Across our simulations, we found that all four robust estimators (L1, S, M, and MM) were less sensitive to model violations in the form of instantaneous evolutionary shifts than the classical L2 estimator (Fig. 3), though this result was primarily restricted to PIC regression (Figs. 3a, c, and e). Specifically, as we increased the severity of model violation in our simulations by increasing shift variance s, the false positive rate of the L2 estimator with PIC regression increased, whereas the robust estimators displayed improved and varying levels of resistance (Fig. 3a). In particular, S and MM estimators were the most resistant to high values of s, whereas the M and L1 estimators exhibited slightly higher false positive rates, but nonetheless fared better than the L2 estimator (Fig. 3a). The mean *P*-value across replicates tracked these differences between estimators as the amount of model violation increased in the simulations (Fig. 3c). Estimates of the slope parameter 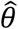 appeared to be unbiased for these estimators, but with higher variance as s increased in our simulations (Fig. 3e). Increasing the number of species underscored the benefits of the four robust estimators over the standard L2 estimator with PIC (Fig. S1), but again not with PGLS (Fig. S2) regression. In all cases, our results corroborate those of Uyeda et al. (2018), who found that classical L2-based regression is highly sensitive to instantaneous evolutionary shifts.

**FIGURE 3.**
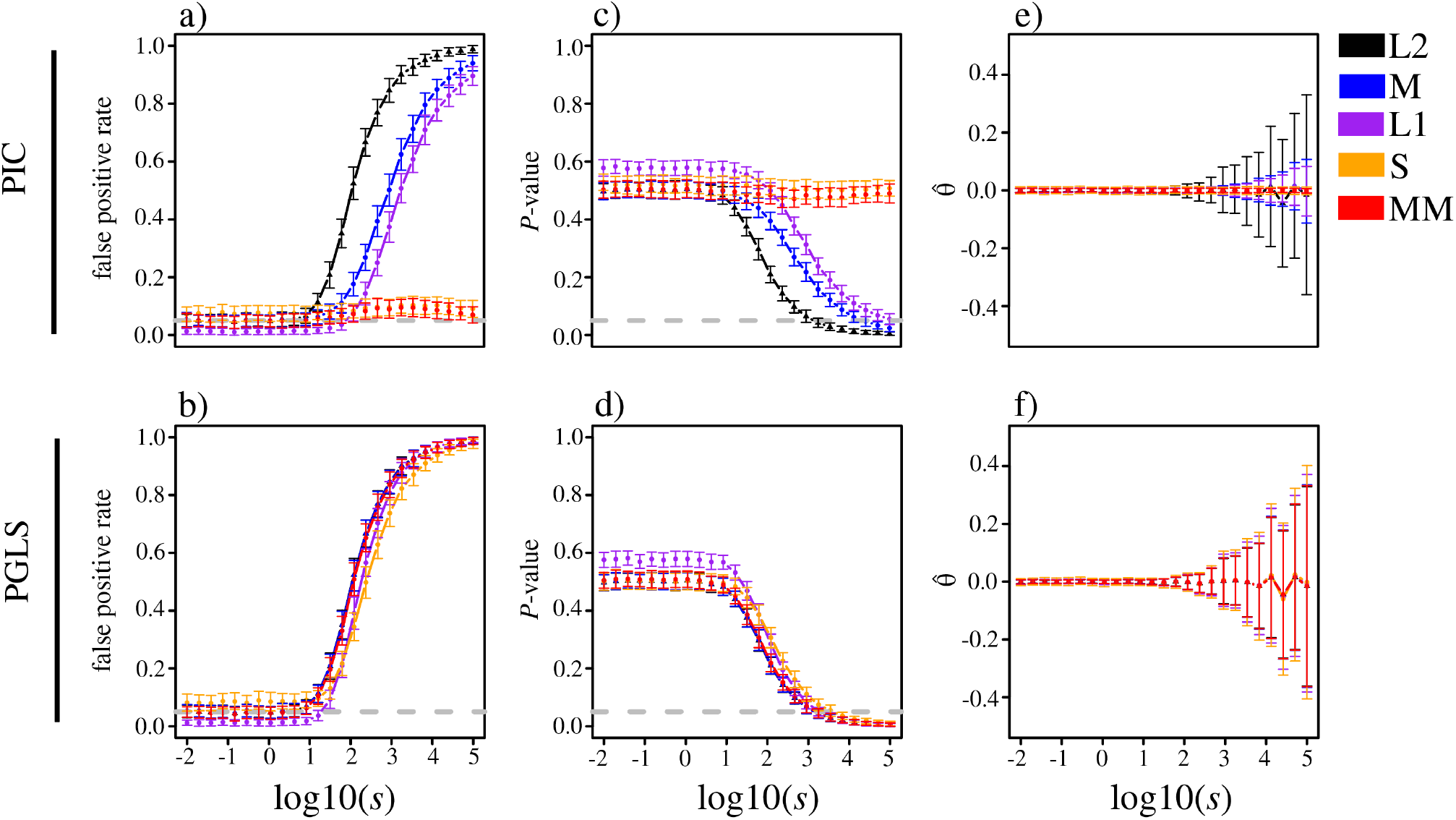
Investigating the impacts of evolutionary model violations for uncorrelated traits simulated under a shift model with variance s in *m* = 256 species. Depicted are the false positive rate measured as the proportion of replicates with *P* ≤ 0.05 (a and b), mean *P*-value (c and d), and estimate 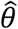 of the slope coefficient *θ* (e and f) plotted as a function of s for each of the five linear estimators (L2, M, L1, S, and MM) with PIC (top) and PGLS (bottom) regression, respectively.

The major advantage of employing robust estimators, particularly S and MM, with PIC regression is summarized by alluvial plots showing their relative fractions of true negatives (*P* > 0.05) and false positives (*P* ≤ 0.05) across simulated replicates (Fig. 4). For simulations with no instantaneous shifts (*s* = 0), and hence no model violations, the L2 and all four robust estimators exhibited low false positive rates (Fig. 4a). In contrast, for simulations with strong instantaneous shifts (*s* = 10^5^), all estimators had high false positive rates except for the S and MM estimators, which controlled the false positive rates (Fig. 4b). Our results suggest that in the presence of uncorrelated evolution between traits and uncertainty in their past dynamics, S and MM estimators with PIC regression represent appropriate statistical models for the PCM toolkit.

**FIGURE 4.**
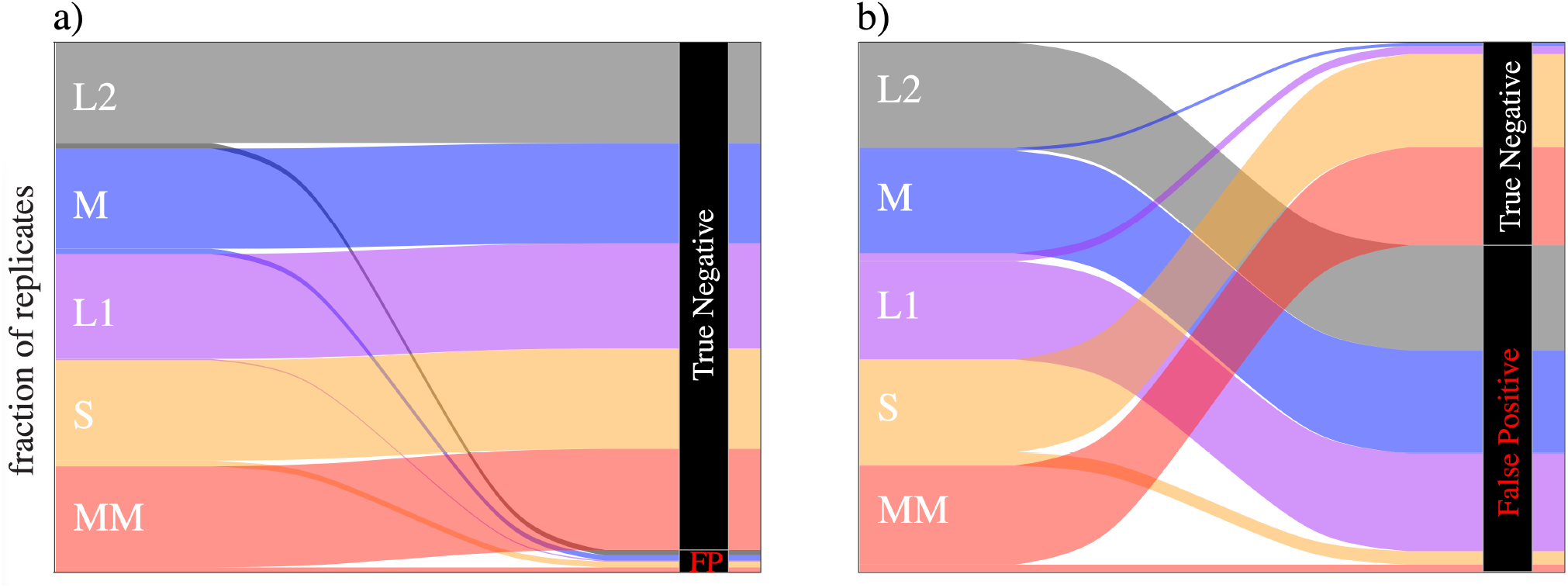
Alluvial plots showing the relative fractions of replicates resulting in either a true negative (*P* > 0.05) or false positive (*P* ≤ 0.05) for applications of the five linear estimators (L2, M, L1, S, and MM) with PIC regression to simulations without any model violations (a; *s* = 0) and with strong model violations (b; *s* = 10^5^).

### Robust Regression Can Identify Correlated Trait Evolution

In addition to assaying robustness to model violations, we evaluated statistical power (true positive rate) of the five estimators (L2, L1, S, M, and MM) to detect true trait relationships (i.e., *ρ* > 0) under the coevo model with PIC and PGLS regression (Fig. 5). Statistical power was highest and comparable with the L2, M, and MM estimators, intermediate with the L1 estimator, and lowest with the S estimator for both PIC (Fig. 5a) and PGLS (Fig. 5b) regression. Unlike our sensitivity to outlier analyses under the shift model (Fig. 4), we did not find major differences in statistical power of the five estimators when comparing PIC to PGLS regression (Figs. 5a vs. 5b), though the S estimator demonstrated slightly higher power with PGLS regression. As expected, power to detect true trait coevolution increased as *ρ* increased, with the S estimator achieving lower power relative to other estimators for moderate to high values of *ρ*. Mimicking our power analyses, mean *P*-value decreased with increasing *ρ* for all estimators, falling below 0.05 for *ρ* > 0.4 with L2, M, and MM estimators, for *ρ* > 0.5 with the L1 estimator, and for *ρ* > 0.7 and 0.9 with the S estimator for PGLS and PIC regression, respectively (Figs. 5c and d). Similar to our results under the shift model, the coevo estimates of the slope coefficient 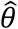 were unbiased, such that the estimates 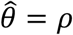 were on average equal to the true values of trait coevolution for both PIC (Fig. 5e) and PGLS (Fig. 5f) regression. Moreover, variability surrounding the estimates 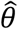 tended to be small, regardless of the true value of trait coevolution *ρ* (Figs. 5e and 5f). Further, all performance results exhibited by the five estimators under the coevo model held across a wide range of sample sizes (Figs. S3 and S4).

**FIGURE 5.**
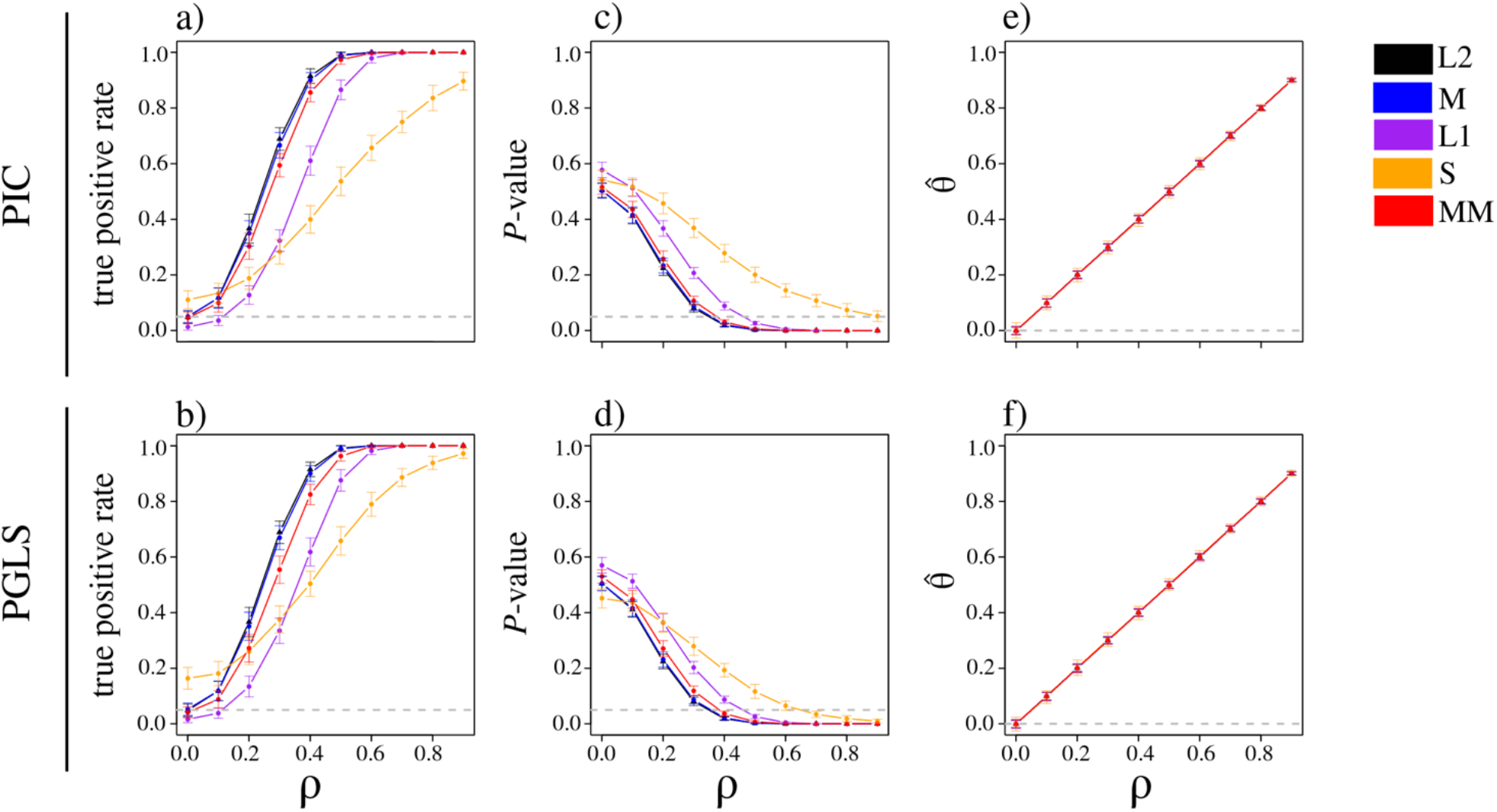
Investigating the impacts of the degree of true trait association for correlated traits simulated with among-trait covariance *ρ* in *m* = 256 species. Depicted are the true positive rate measured as the proportion of replicates with *P* ≤ 0.05 (a and b), mean *P*-value (c and d), and estimate 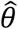 of the slope coefficient *θ* (e and f) plotted as a function of *ρ* for each of the five linear estimators (L2, M, L1, S, and MM) with PIC (top) and PGLS (bottom) regression.

To better understand the impact of *ρ* on the power to detect true trait coevolution, we generated alluvial plots depicting the relative fractions of false negatives (*P* > 0.05) and true positives (*P* ≤ 0.05) for applications of the five linear estimators with PIC regression to simulated replicates with true evolutionary covariances of *ρ* = 0.3, 0.6, and 0.9 (Fig. 6). When true evolutionary covariance was weak (*ρ* = 0.3), power was less than 0.5 for all methods, with the L1 and S estimators demonstrating the lowest powers (Fig. 6a). In contrast, when evolutionary covariance was moderately strong (*ρ* = 0.6), all methods had powers of 1.0 or close to 1.0, except for the S estimator, which displayed a false negative rate of 0.34 (Fig. 6b). Finally, in the extreme setting of strong evolutionary covariance (*ρ* = 0.9), all estimators had high power, with each achieving a power of 1.0 except for the S estimator, for which the false negative rate decreased to 0.10 (Fig. 6c). Taken together, results from the shift and coevo models suggest that application of the MM estimator with PIC regression yields both the greatest robustness to false signatures of trait coevolution and the highest power to detect true trait coevolution.

**FIGURE 6.**
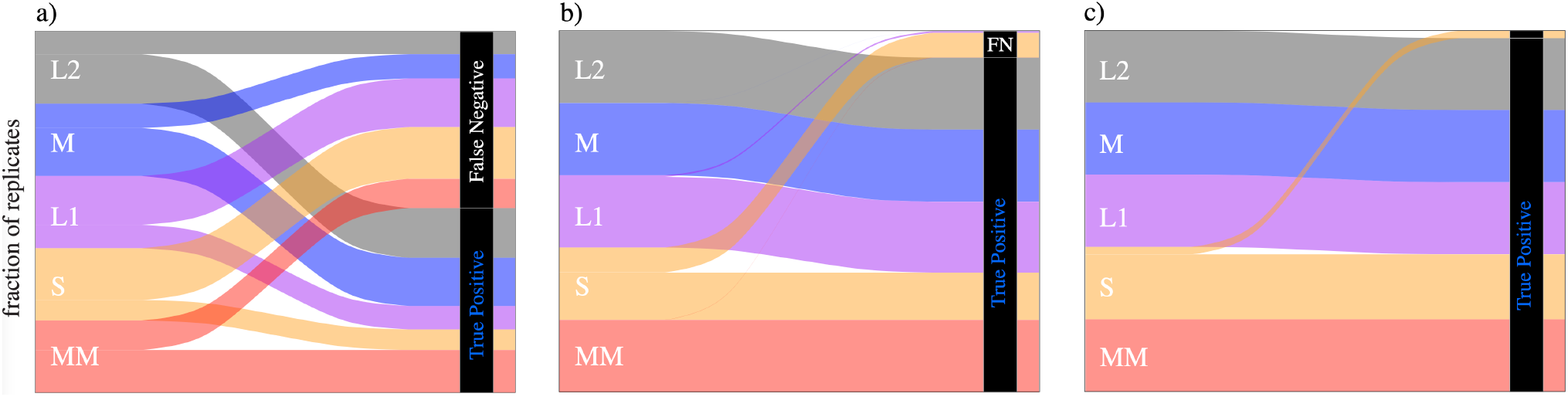
Alluvial plots showing the relative fractions of replicates resulting in either a true positive (*P* ≤ 0.05) or false negative (*P* > 0.05) for applications of the five linear estimators (L2, M, L1, S, and MM) with PIC regression to simulations with true evolutionary covariance *ρ* = 0.3 (a), *ρ* = 0.6 (b), and *ρ* = 0.9 (c).

### Application of Robust Regression to Empirical Gene Expression Data

Our simulation experiments highlighted the varying robustness and statistical power of the five estimators (L2, M, L1, S, and MM) across an array of scenarios with and without trait coevolution. Hence, we next explored whether application of these estimators to an empirical dataset would yield different conclusions about trait evolution. To this end, we examined the agreement between predictions employing robust estimators and the classical L2 estimator with PIC regression applied to a dataset composing expression measurements of 5,615 genes in 11 tissues from nine species (Brawand et al. 2011; see Methods for details). We chose this dataset because adaptation often proceeds through regulatory modifications that alter spatial or temporal expression levels of genes (King and Wilson 1975; Wray et al. 2003; Carroll 2005; Jones et al. 2012; Mack et al. 2018), with recent studies showing that such changes can occur as rapid evolutionary shifts (Barua and Mikheyev 2020; Hamann et al, 2021). Thus, these spatial gene expression data represent an excellent setting to showcase the empirical performance of robust estimators.

We first investigated whether there were global differences in the abilities of the five estimators to detect correlated expression evolution by assaying relationships between the expression measurements of genes in the same tissue in females and males (Fig. S5). Indeed, distributions of *P*-values varied across estimators, and were smallest with the S estimator, then with MM, M, L2, and finally L1 estimators (Figs. S5a-c). Because this coarse overview of *P*-value distributions across estimators ignores information about how often the *P*-value differs for one estimator when controlling for the same tested gene, we also computed differences in gene-specific *P*-values between each of the robust estimators and the L2 estimator. Comparison of the resulting distributions among estimators revealed that *P*-value differences with the L2 estimator were smallest and least variable for the M estimator, and largest and most variable for the L1 estimator (Figs. S5d-f). Moreover, relative to the L2 estimator, *P*-values were smaller for the L1 estimator and larger for the S and MM estimators (Figs. S5d-f). Similarly, we calculated differences in estimates of the slope, which measures the magnitude and direction of the expression relationship, between each of the robust estimators and the L2 estimator. Comparison of these distributions demonstrated that slope differences with the L2 estimator were small for all robust estimators on average, though tended to be least variable for M estimation (Figs. S5g-i).

To more finely examine alignment between the four robust estimators and the L2 estimator, we compared the proportion of genes at which a given robust estimator agrees or disagrees with the L2 estimator rejecting (*P* ≤ 0.05) or not rejecting (*P* > 0.05) the null hypothesis of no coevolution between female and male expression in heart, kidney, and brain (Fig. 7). As expected from our broader analysis (Fig. S5), the M estimator tended to agree with the L2 estimator, with a skew toward the M estimator rejecting the null hypothesis when the L2 estimator does not reject it more often than not rejecting the null hypothesis when the L2 estimator rejects it (Fig. 7a). The behavior of the L1 estimator was similar (Fig. 7b), whereas the S and MM estimators demonstrated the same skew but also often did not agree with the L2 estimator (Figs. 7c and d).

**FIGURE 7.**
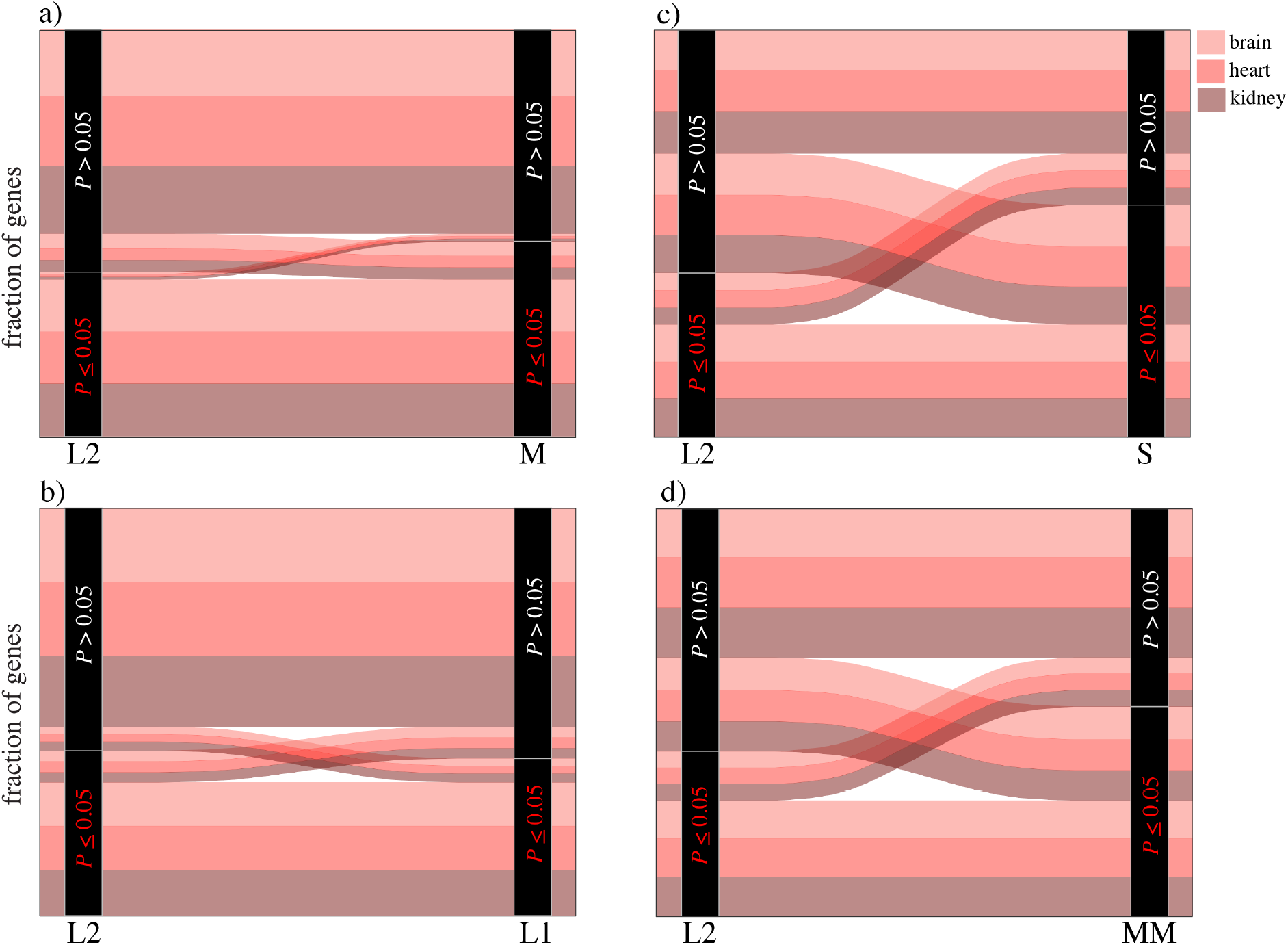
Alluvial plots showing relative fractions of genes with significant (*P* ≤ 0.05) and nonsignificant (*P* > 0.05) correlated expression in female and male brain, heart, and kidney tissues from nine species. Depicted are inferences from the standard L2 estimator (left side of each panel) compared with M (a), L1 (b), S (c), and MM (d) estimators with PIC regression. For example, in panel (a), the fraction of non-significant genes with both L2 and M estimation are shown at the top, while significant genes with L2 and non-significant genes with M estimation result in mismatched connections.

As a final investigation into the differences between the robust and classical L2 estimators, we performed case studies of genes for which robust estimators disagreed with the L2 estimator in each of the three tissues. We first considered one gene in each tissue for which the null hypothesis of no expression coevolution was rejected by the L2 estimator, but not by at least one robust estimator (Figs. 8a-c). In such cases, the L2 estimator was potentially misled into identifying gene expression coevolution by a single high-leverage outlier (Figs. 8a and c). As expected by the high level of disagreement observed in our alluvial plots (Fig. 7), both S and MM estimators did not reject the null hypothesis when the L2 estimator rejected the null hypothesis for female and male coevolution of *TP53I11* (ENSG00000175274) in heart (Fig. 8a), *NAV1* (ENSG00000134369) in kidney (Fig. 8b), and *DIPK1B* (ENSG00000165716) in brain (Fig. 8c). *TP53I11* encodes a protein predicted to negatively regulate cell population proliferation (The Alliance of Genome Resources Consortium 2019), *NAV1* encodes a protein hypothesized to be involved in neuronal development and regeneration (O’Leary et al. 2016), and *DIPK1B* encodes a transmembrane protein whose function is unknown (O’Leary et al. 2016). Thus, there does not appear to be a clear biological reason for coevolution of any of these genes between females and males in the tissues in which they were uncovered by L2-based PIC regression.

**FIGURE 8.**
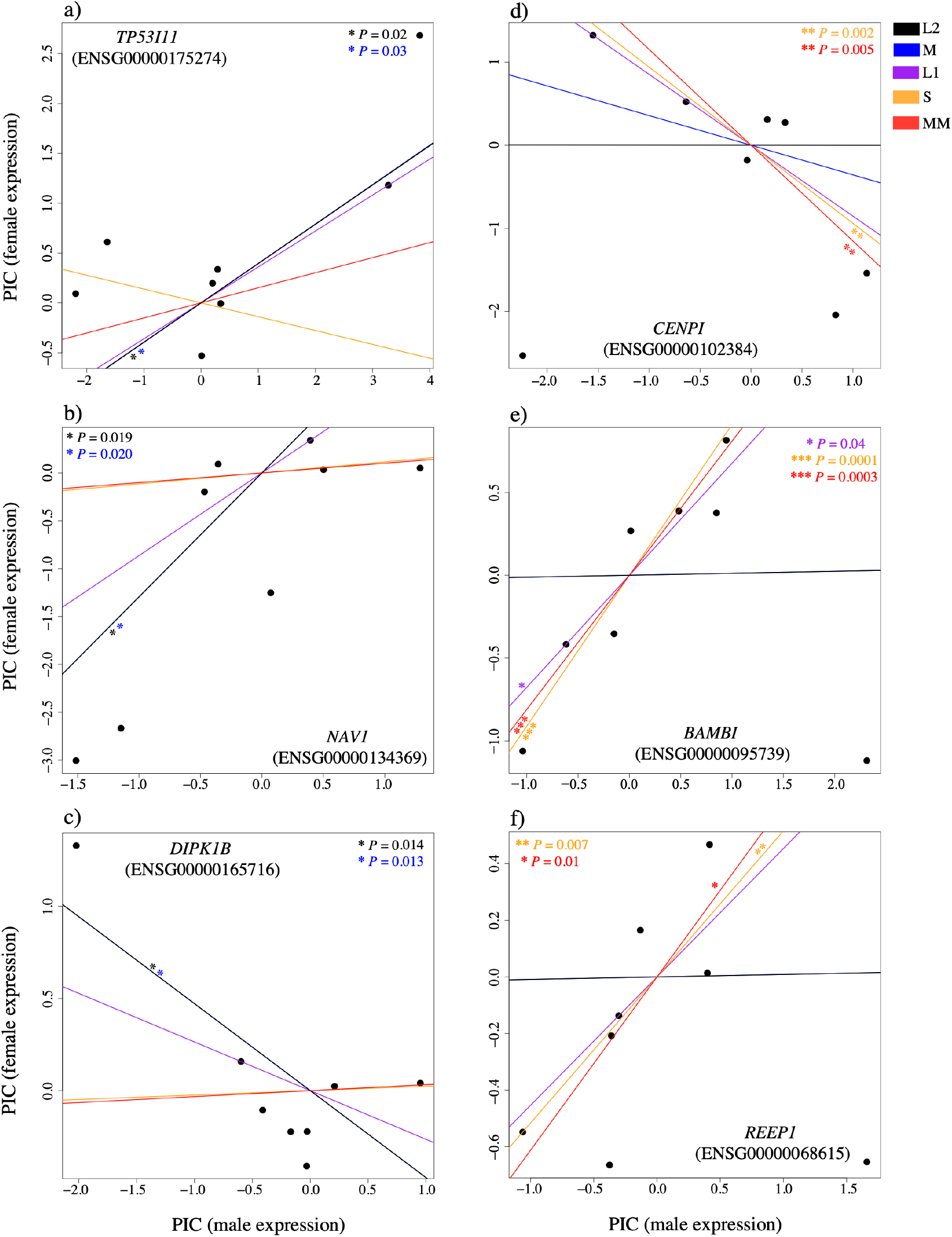
Examples highlighting differences between the L2 estimator and four robust estimators with PIC regression for testing relationships between expression in female and male heart (a and d), kidney (b and e), and brain (c and f) tissues.

Next, we examined one gene in each tissue for which the null hypothesis of no expression coevolution was rejected by at least one robust estimator, but not by the L2 estimator (Figs. 8d-f). Again, in all cases, the L2 estimator appeared to be swayed by a high-leverage outlier. Specifically, the L2 estimator did not reject the null hypothesis, whereas both S and MM estimators provided evidence of a significant relationship (*P* ≤ 0.05) for female and male coevolution of three genes: *CENPI* (ENSG00000102384) in heart (Fig. 8d), *BAMBI* (ENSG00000095739) in kidney (Fig. 8e), and *REEP1* (ENSG00000068615) in brain (Fig. 8f). Intriguingly, *CENPI* encodes a key mitotic protein involved in gonad development and gametogenesis (O’Leary et al. 2016), which may provide a basis for coevolution of its expression between female and male tissues. *BAMBI* encodes a transmembrane glycoprotein that limits signaling of the TGF-beta gene family during early embryogenesis (O’Leary et al. 2016) and is associated with diabetes insipidus and childhood-onset diabetes mellitus (Awata et al. 2000; El-Shanti et al. 2000), perhaps explaining its coevolution in female and male kidney. *REEP1* encodes a mitochondrial protein that enhances the cell surface expression of odorant receptors (O’Leary et al. 2016) and is associated with several neurodegenerative diseases (Züchner et al. 2006; Beetz et al. 2012; Kanwal et al. 2021), which may account for its coevolution in female and male brain. Thus, all genes appear to have some biological support for either their sex-specific or tissue-specific covariation.

## Discussion

Pseudoreplication due to shared ancestry is a bane of comparative biologists that can confound inferences of trait coevolution. Robust phylogenetic regression provides a solution to this problem by building a better defense against model violations and statistical outliers commonly encountered in trait data. Here we sought to build upon the PCM framework by introducing and exploring the application of a suite of new robust linear estimators for phylogenetic regression. Our analyses revealed that these robust estimators hold up to their namesake in both Felsenstein’s (1985) worst-case scenario and a simple bifurcating tree (Fig. 2). Despite deliberate use of challenging evolutionary and statistical scenarios (Uyeda et al. 2018), several robust estimators demonstrated improvements over the classical L2 estimators in many contexts (Figs. 3–6 and S1-S4), with particularly favorable performance observed for the S and MM estimators with PIC regression (Figs. 3–6, S1, and S3). Specifically, these estimators tended to not be misled by model violations and outliers into detecting false signals of trait associations (Figs. 3, 4, and S1), and also demonstrated high statistical power for detecting true trait associations (Figs. 5, 6, S3, and S4). In contrast, and in strong agreement with previous studies (e.g., Uyeda et al. 2018), we find that classical L2-based phylogenetic regression, which has long remained the *de facto* approach for testing trait coevolution, suffers high systematic error in many scenarios.

One pervasive trend in our analyses was the contrasting robustness of phylogenetic regression based on PIC and PGLS principles (Figs. 2, 3, S1, and S2). PIC and PGLS regression are mathematically equivalent for L2 estimation (Blomberg et al. 2012), and yet, the performance of robust estimators differed markedly between the two approaches, particularly in the presence of outliers (Figs. 3, S1, and S2). While we observed considerable advantages in using robust estimators with PIC regression (e.g., Fig. 3a), these benefits were less apparent with PGLS regression (e.g., Fig. 3b). There may be several reasons for these discrepancies, and our analyses suggest that the manner by which traits are projected may hold clues. Our simple demonstration of phylogenetic regression in Figure 2 offers insights into differences when using robust estimators with PGLS (Figs. 2c and d) verses PIC (Figs. 2e and f). By transforming the traits into a set of independent contrasts, PIC regression has effectively one fewer input observation (i.e., *n* = *m* – 1 contrasts), which is predicted to reduce significance when testing a null hypothesis by increasing the standard error of the test statistic. Importantly, these contrasts collectively express all variation present among a set of *m* species related by a phylogeny (Felsenstein 1985), and by doing so, encode information about ancestral traits and the phylogenetic locations of contrasts that may represent outliers (e.g., dark points in Figs. 2e and f). Thus, clustering of PICs tended to isolate the single outlying contrast representing the specific branch location of the model violation itself (e.g., marked dashes in Figs. 2a and b correspond to dark points in Figs. 2e and 2f). In contrast, trait projections based on the PGLS transformation yielded two large partitions corresponding to two clusters: species that did (dark points in Figs. 2c and d) and that did not (white points in Figs. 2c and d) experience the model violation in their phylogenetic history. That is, PGLS transformations separate the trait values into two clusters as a simple binary response to the presence of the model violation in the lineage history of a set of species, whereas PIC effectively distinguishes the presence of a single distinctive outlier (Fig. 2c vs. Fig. 2e). PIC regression therefore allowed the robust estimators to take full advantage of the information present within trait contrasts, improving resistance to model violations; this can be observed when comparing estimated relationships of L2 with M, L1, S, and MM estimators in Figure 2.

Throughout our analyses, perhaps most apparent were the advantages of using the MM estimator with PIC regression, which achieved the highest observed resistance to evolutionary outliers (Figs. 3 and S1) while retaining sufficient power for detecting true trait associations (Figs. 5 and S3). Specifically, whereas MM-based PIC regression was largely unaffected by the presence of very large magnitude model violations in extreme evolutionary scenarios (e.g., Fig. 3a), it displayed comparable power to classical L2-based PIC regression for detecting true trait associations (Fig. 5). Thus, application of MM-based PIC regression may serve as a single best general-purpose strategy for assaying trait associations. Further, among the four robust estimators explored here (M, L1, S, and MM), the L1 estimator appeared to be the least robust, and yet still outperformed the L2 estimator in many cases (Figs. 3 and S1). Collectively, these findings underscore improvements of robust phylogenetic regression compared to classical L2-based approaches, demonstrating that robust estimators may be used *in lieu* of the classical L2 estimator to provide more reliable inferences of trait coevolution. In this sense, robust regression represents a much-needed answer to ongoing questions and debates about coevolution inferences in the presence of evolutionary model violations.

Our empirical studies of gene expression in three female and male tissues also provided support for improved inferences with robust estimators compared to standard L2-based phylogenetic regression. In particular, we showed that there were often differences between genes inferred as coevolving in female and male tissues by robust and the L2 estimator (Figs. S5 and 7), and that such differences may be indicative of their varying sensitivities to outliers (Figs. 8a-c) or powers for detecting true coevolutionary relationships (Figs. 8d-f). For example, evaluation of female and male coevolution in heart tissue resulted in rejection of the null hypothesis by the L2 estimator but not by either S or MM estimators for the *TP53I11* gene (Fig. 8a), and conversely, in rejection of the null hypothesis by both S and MM estimators but not by the L2 estimator for the *CENPI* gene (Fig. 8d). In both cases, these differences were driven by a single outlier that either reduced the robustness (Fig. 8a) or power (Fig. 8d) of the L2 estimator, highlighting the sensitivity of classical L2-based phylogenetic regression to model violations and underscoring the benefits of robust estimators in this context. Consistent with this observation, *TP53I11* does not appear to be related to sex or heart tissue, whereas *CENPI* is involved in sex-specific functions (O’Leary et al. 2016) that may help explain its apparent coevolution in female and male tissues. Hence, this example highlights the utility of robust phylogenetic regression in detecting biologically relevant trait associations when classical methods may fail in the presence of model violations. When considering the implications of these findings, we found the following quote from Tukey (1975) to be quite fitting: “It is perfectly proper to use both classical and robust/resistance methods routinely, and only worry when they differ enough to matter. But when they differ, you should think hard.” Thus, contrasting the results of different linear estimators (both robust and classical) may be fruitful if trait data are suspect of model violations.

Though robust phylogenetic regression is a rigorous and powerful solution to the problem of evolutionary pseudoreplication, we neither claim that it is the only one, nor that it solves this problem completely. However, our results do suggest that robust regression is likely to be helpful in many scenarios of unreplicated evolutionary events, and therefore represents a step toward a common solution. Uyeda et al. (2018) argued for a philosophical unification of phylogenetic natural history and *a priori* hypothesis testing (e.g., Maddison et al. 2007; Alfaro et al. 2009; Stadler 2011; FitzJohn 2012; Rabosky 2014); because robust regression decreases sensitivity to phylogenetic outliers, this approach may help achieve both goals by accounting for hidden unreplicated evolutionary events in the tree when testing for trait associations. Specifically, large differences between classical L2 phylogenetic regression and robust methods may be indicative of interesting evolutionary phenomena, represented by high-leverage outliers within the ancestral history of a focal clade of interest (e.g., Fig. 2). Whereas we examined the performance of robust phylogenetic regression in a few key settings, future investigations will be necessary to further our understanding of its sensitivity and power in the presence of a diversity of model violations and evolutionary scenarios. In particular, phylogenetic regression is typically conducted under the assumption of BM evolution (or similar models) in the absence of shifts, and thus, recent contributions to multivariate models of trait coevolution and shifts that include both between-species correlations (covariance due to shared species histories) and between-trait correlations (covariance due to shared trait functions) hold particular promise (Uyeda and Harmon 2014; Bastide et al. 2017, 2018a; Duchen et al. 2017). Strategies such as Bayesian mixture models (Uyeda and Harmon 2014; Uyeda et al. 2017, 2018) and the implementation of heavy-tailed distributions for modeling error terms while fitting regression models (Landis et al. 2013; Elliot and Mooers 2014) have also been suggested to deal with these issues (Uyeda et al. 2018), as well as goodness-of-fit tests and graphical models (Höhna et al. 2014, 2016). Future work will be useful for evaluating different types of model violations and scenarios of non-BM evolution, non-bifurcating trees (Bastide et al. 2018b), and more complex evolutionary shifts (Duchen et al. 2017; Bastide et al. 2018a; Mitov et al. 2019). Collectively, our findings join arguments for increased vigilance against evolutionary pseudoreplication and a better understanding of phylogenetic modeling assumptions (i.e., Revell 2010; Ho and Ané 2014; Mundry 2014; Maddison and FitzJohn 2015; Uyeda et al. 2018; Ives 2019), setting the stage for a shift in method development in this important area.

## Software Availability

We have implemented a suite of functions for conducting robust phylogenetic regression in the R package ROBRT (ROBust Regression on Trees). ROBRT includes functions for computing each of the four robust estimators explored in this study (M, L1, S, and MM), as well as the standard L2 estimator. ROBRT is freely available for open use by the community (https://github.com/radamsRHA/ROBRT), and requires the dependencies APE (Paradis et al. 2004) and PHYTOOLS (Revell 2012).

## Funding and Acknowledgements

This work was supported by National Science Foundation grants DEB-1949268, DEB-2001059, DBI-2130666, and BCS-2001063, the National Institutes of Health grants R35GM142438 and R35GM128590. The authors would like to acknowledge the use of services provided by Research Computing at the Florida Atlantic University.

