## Supplementary Figures for "Robust phylogenetic regression"

### SUPPLEMENTARY MATERIALS

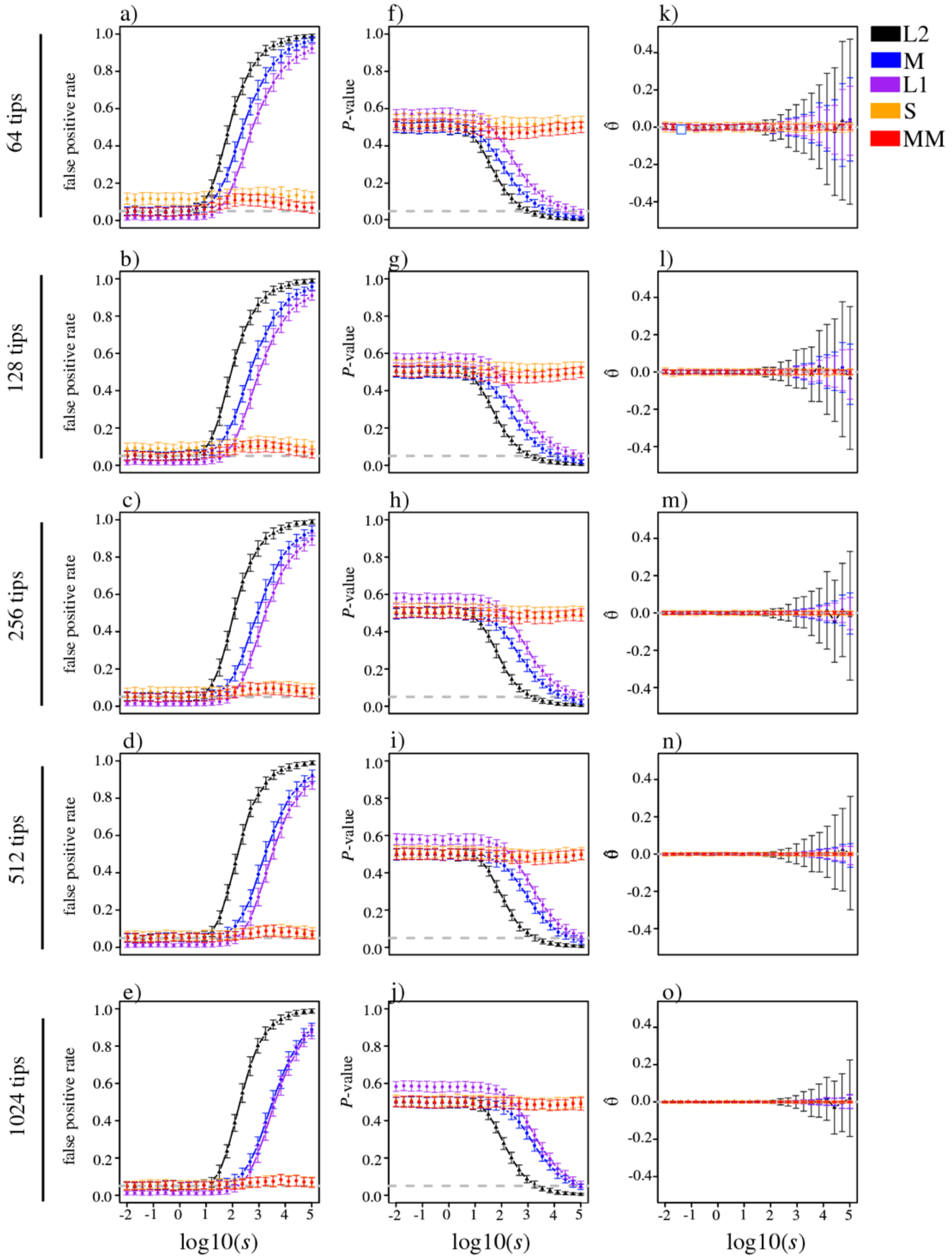


SUPPLEMENTARY FIGURE 1. Investigating the impacts of evolutionary model violations for uncorrelated traits simulated under a shift model with variance $s$. Depicted are the false positive rate measured as the proportion of replicates with $P \leq 0.05$ (a-e), mean $P$-value (f-j), and estimate $\hat{\theta}$ of the slope coefficient $\theta$ (k-o) plotted as a function of $s$ for each of the five linear estimators (L2, M, L1, S, and MM) with PIC regression.

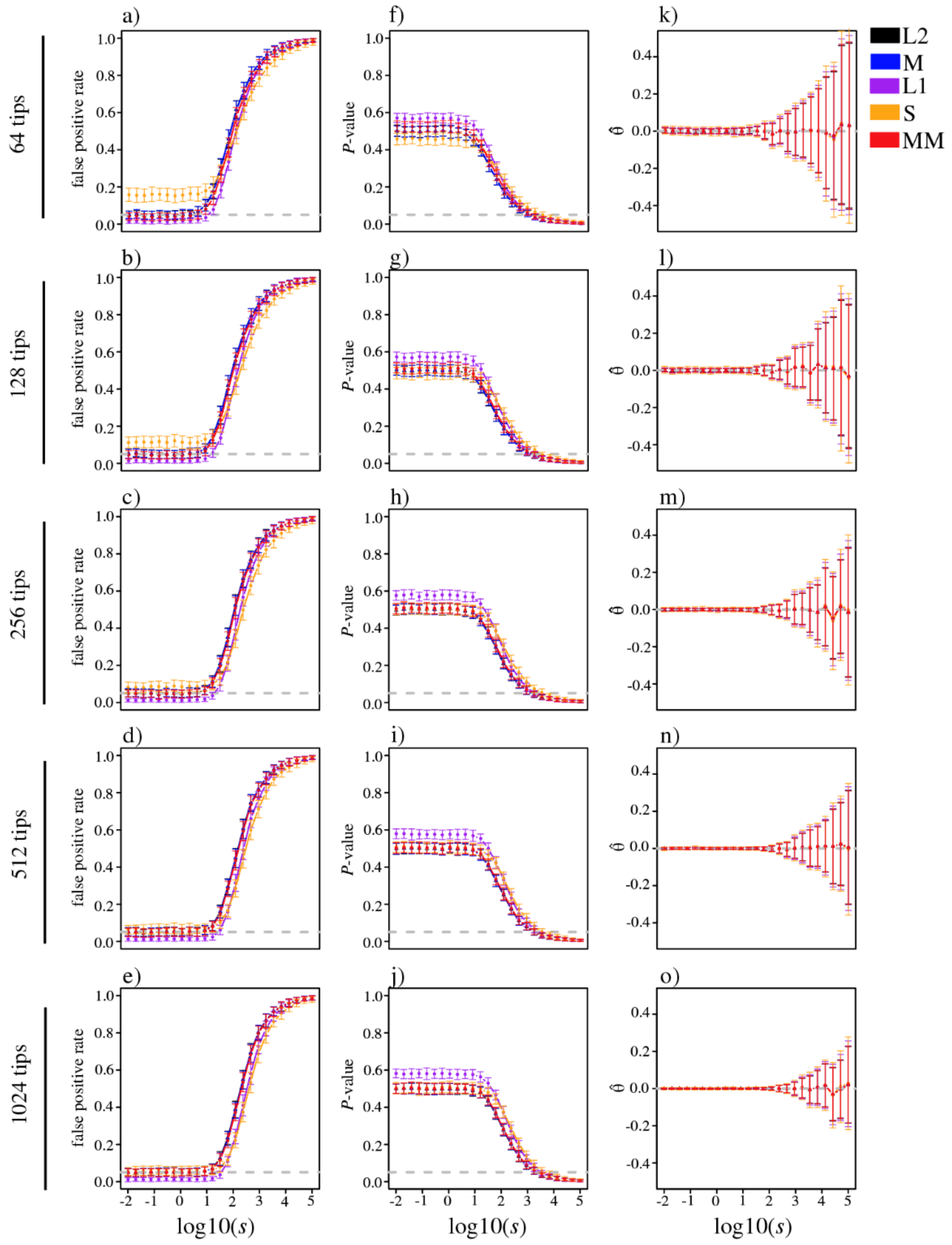


SUPPLEMENTARY FIGURE 2. Investigating the impacts of evolutionary model violations for uncorrelated traits simulated under a shift model with variance $s$. Depicted are the false positive rate measured as the proportion of replicates with $P \leq 0.05$ (a-e), mean $P$-value (f-j), and estimate $\hat{\theta}$ of the slope coefficient $\theta$ (k-o) plotted as a function of $s$ for each of the five linear estimators (L2, M, L1, S, and MM) with PGLS regression.

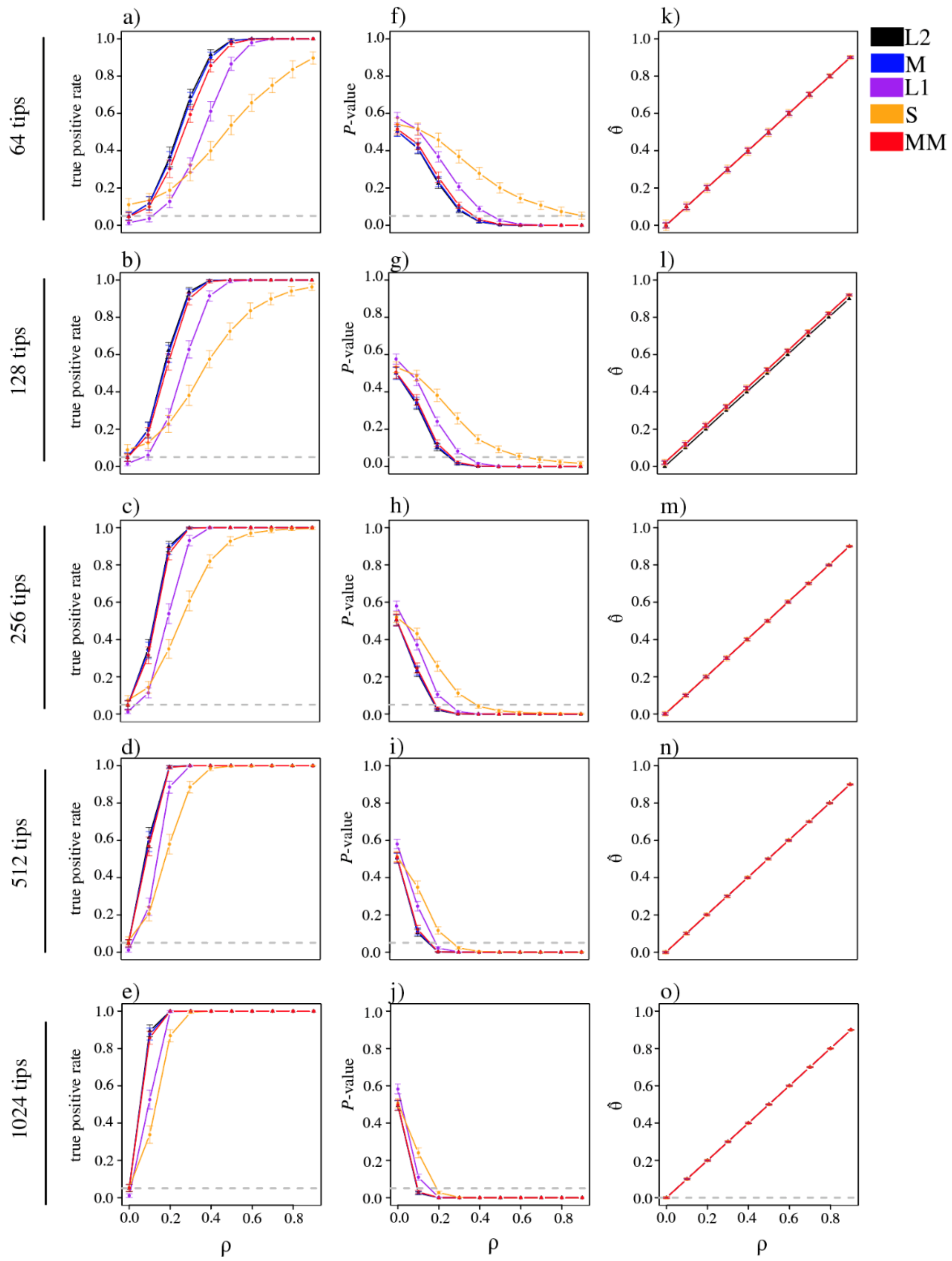


SUPPLEMENTARY FIGURE 3. Investigating the impacts of the degree of true trait association for correlated traits simulated with true trait covariance $\rho$. Depicted are the true positive rate measured as the proportion of replicates with $P \leq 0.05$ (a-e), mean $P$-value (f-j), and estimate $\hat{\theta}$ of the slope coefficient $\theta$ (k-o) plotted as a function of $\rho$ for each of the five linear estimators (L2, M, L1, S, and MM) with PIC regression.

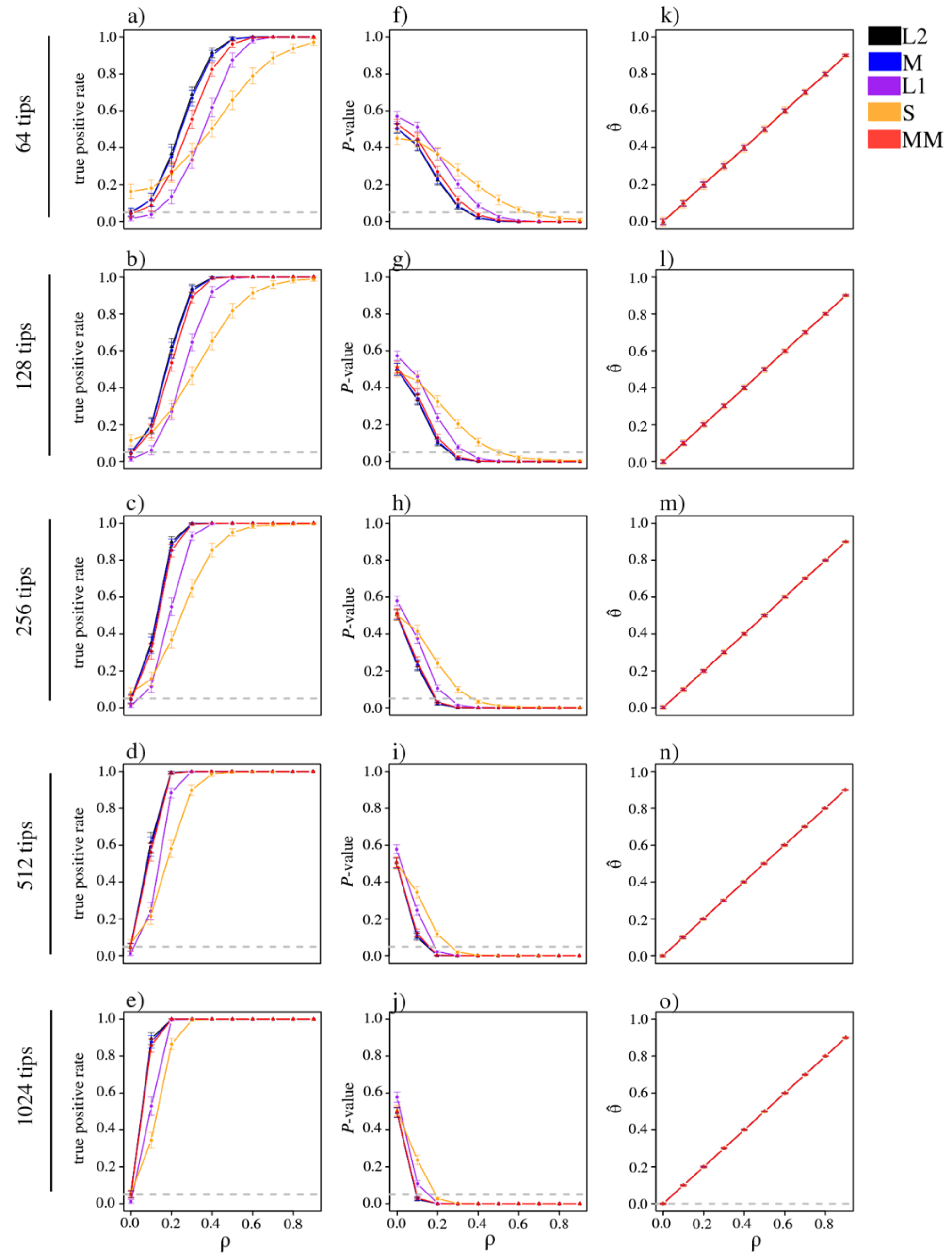


SUPPLEMENTARY FIGURE 4. Investigating the impacts of the degree of true trait association for correlated traits simulated with true trait covariance $\rho$. Depicted are the true positive rate measured as the proportion of replicates with $P \leq 0.05$ (a-e), mean $P$-value (f-j), and estimate $\hat{\theta}$ of the slope coeffecient $\theta$ (k-o) plotted as a function of $\rho$ for each of the five linear estimators (L2, M, L1, S, and MM) with PGLS regression.

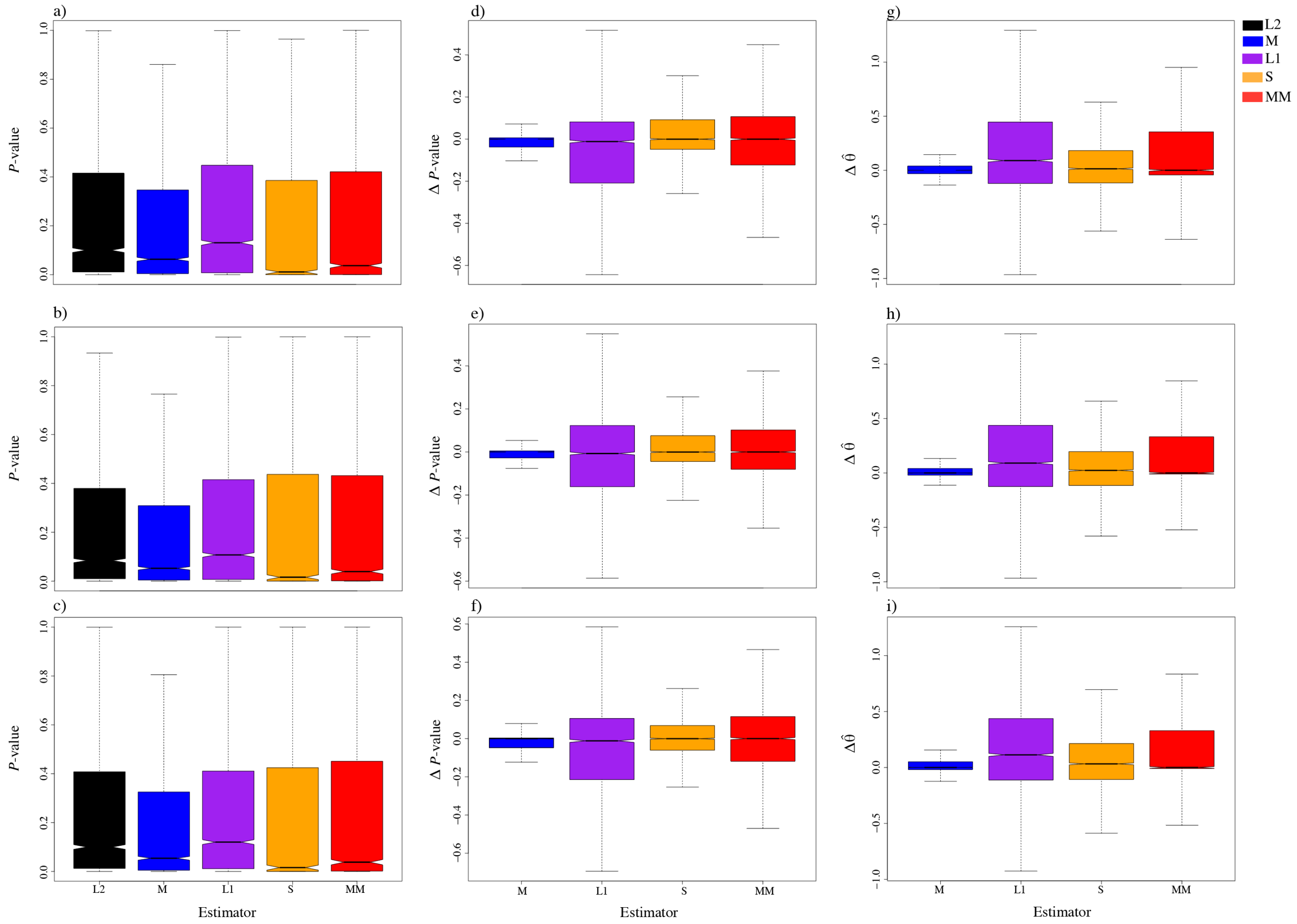


SUPPLEMENTARY FIGURE 5. Exploring robust regression for predicting female expression as a function of male expression in three tissues. Distributions of *P*-value for each of the five estimators (L2, M, L1, S, and MM; a-c), Δ*P*-value (difference between *P*-values of each robust estimator and the L2 estimator; d-f), and $\Delta\hat{\theta}$ (difference between coefficient estimates of each robust estimator and the L2 estimator; g-i) with PIC regression applied to expression data from 5,615 genes in heart (top row), kidney (center row), and brain (bottom row) tissues. Outliers have been removed from boxplots to better visualize differences among distributions.
